# The evolution of host exploitation by parasitoids: the timing of attack and consumption

**DOI:** 10.1101/2025.02.22.639661

**Authors:** Ryuichiro Isshiki, Ryosuke Iritani

## Abstract

Parasitoid wasps exhibit remarkable diversity in life-history traits and are categorized into two major groups based on the timing of emergence: those that begin consuming host tissues immediately after hatching (“idiobiont”) and those that can delay the emergence depending on host maturation (“koinobiont”). Although delayed emergence allows parasitoids to exploit unparasitized, young hosts, it incurs a mortality cost during the waiting period. While numerous empirical studies have examined the adaptive significance of this trait, how host stage-structure and ecological factors jointly shape the evolution of emergence timing remains poorly understood in a formalized context. Here, we develop a stage-structured mathematical model to analyze the evolutionary dynamics of delayed emergence and its consequences for host exploitation. By explicitly incorporating host developmental stages, our model explores the joint evolution of two traits: the preference for young versus old host, and the timing of emergence. Our results reveal that the optimal strategy is determined by the relative reproductive value of host stages, which depends on the balance between host growth and mortality. We demonstrate that delayed emergence evolves when the future value of a growing host outweighs the immediate cost of mortality, thereby driving the divergence between idiobiont and koinobiont strategies. These findings provide a theoretical framework that unifies the diversity of parasitoid life histories.

**Significance Statement:** Parasitoids exhibit striking diversity in life-history traits such as body size, fecundity, and clutch size, reflecting adaptations to host exploitation. A key behavioral distinction is whether parasitoids consume the host immediately (idiobionts) or can delay emergence while the host continues to grow (koinobionts). Many empirical studies have suggested and tested some hypothesis. On the other hand, the adaptive significance of delayed emergence have not been formulated yet, so the quantitative relationship between selection pressure and life history traits is yet to be better understood. Here, we develop a mathematical model that integrates host growth with parasitoid emergence timing. Our analysis supports the adaptive significance of delayed emergence and shows how delayed emergence can promote diversification of parasitoid life-history strategies. These results suggest that delayed emergence is a key driver of idiobiont–koinobiont dichotomy, providing new insights into the evolutionary basis of parasitoid–host interactions and aligning directly with the central aims of behavioral ecology.

## Introduction

Resource exploitation strategies, specifically their timing and allocation, are fundamental life-history traits. Parasitoids, particularly wasps, serve as an excellent model system for studying life-history traits, with their variations in parasitoids particularly pronounced (Godfray 1994; Mayhew and Blackburn 1999; Jervis et al. 2008; Wajnberg et al. 2008; Jervis and Ferns 2011; Quicke 2014): for example, the order *Hymenoptera* comprises at least 200,000 species by a conservative estimate (Pennacchio and Strand 2006), probably each utilizing different host species (mostly arthropods). Also, wasp species exhibit life-history trait variations, by a factor of 18 in body-size (Jervis et al. 2003), more than one hundred in clutch size and lifetime potential fecundity (Jervis et al. 2008), with these variations presumably resulting from adaptation to host species. Parasitoid life-history traits have significant impacts on numerous processes such as parasitoid-prey interactions (Lafferty et al. 2008), policy making in pest management (Wang et al. 2019), and coevolution (Amoroso and Antonovics 2020). Hence, improving our understanding of the evolution of parasitoids provides crucial implications for biodiversity and its applications.

In parasitoid wasps, life-history trait variations are typically found in sequential timings of host exploitation, from attacking, hatching to emerging (Alonzo and Kindsvater 2008). For example, some species emerge immediately after oviposition (e.g., *Ichneumon promissorius*; Carpenter 1995), while others emerge some time after (e.g., *Venturia canescens*; Beckage & Riddiford 1978; Sequeira and Mackauer, 1992a; Harvey, 1994; Harvey & Vet, 1996; Harvey, 2000; Castelo et al., 2010; Harvey et al., 2012). Such differences in the exploitation timings determine the degree to which parasitoids acquire resources from younger (and thus smaller) versus older (thus larger) hosts, which as a consequence influences the population dynamics of younger and older hosts. In other words, the evolution of exploitation timings strongly influences evolutionary and ecological aspects of parasitoid lives.

The delayed emergence, resulting from parasitoids not consuming their hosts immediately after the attack, is a key factor in creating diversity in the life-history characteristics of parasitoids, including larval mortality, fecundity, and egg size (Jervis and Ferns 2011). Parasitoid wasps can be categorized into two groups according to this trait; namely, “idiophytic” species (‘idio’ means ‘own/individual/personal’ descending from Greek, indicating that hosts are personalized or monopolized by a single parasitoid individual) that consume the host immediately after the attack, and “koinophytic” species (‘koino’ means ‘common’, also from Greek, indicating parasitoids live with a living host) that allow hosts to continue to feed and grow after parasitization (Haeselbarth 1978; Askew &Shaw 1986). By applying this criterion for a wider range of parasitoid wasp species, the major dichotomy is proposed in the larval developmental pattern of parasitoid wasps: idiobiont-koinobiont extremes (Askew&Shaw 1986). The delayed emergence can therefore characterize two major categories of parasitoids (Askew&Shaw 1986; Godfray 1994) with different life-history characteristics. The dichotomy is often used as a basis to understand the life-history traits in parasitoid wasps (Godfray, 1994; Jervis, 2008; Quicke, 2014), although in many koinobiont species, they exhibit delayed emergence with various durations depending on the host stage (Harvey et al., 1994; Harvey et al., 2000; Harvey & Strand, 2002; Harvey et al., 2016).

Studies testing the dichotomous hypothesis examine the relationship between delayed emergence trait and other traits. Statistical analyses that control for phylogenetic correlations also provide some support for the dichotomy hypothesis, suggesting possible mechanisms for the observed pattern of association between life-history traits (Mayhew and Blackburn 1999). For example, koinobionts exhibit higher larval mortality, greater fecundity, and smaller egg sizes compared to idiobionts (Mayhew and Blackburn 1999). Mayhew and Blackburn (1999) suggest that these trait associations are explained by both selective pressures and developmental constraints: the higher mortality rate may result from the increased exposure to predators or immaturity of the host. This in turn favours higher fecundity but at the cost of smaller egg sizes (Mayhew and Blackburn 1999). Furthermore, idiobionts require a substantial reproductive investment for egg production because females must package all the nutrients – primarily proteins – necessary for the completion of embryogenesis into these eggs (Jervis & Kidd, 1986). Consequently, protein-rich “anhydropic” eggs have evolved. Due to the large size of these eggs, most idiobiont females cannot carry many eggs at one time and the fecundity tends to be low (Jervis & Kidd, 1986). In contrast, “hydropic” eggs that contain little protein have evolved in koinobionts, likely resulting from the evolution of an extra-embryonic membrane that enables the embryo to uptake proteins and other nutrients directly from the host haemolymph (Jervis & Kidd, 1986). This adaptation allows koinobiont females to produce many small eggs and the handling time for oviposition is often shorter than idiobiont’s (Jervis & Kidd, 1986). Therefore, various life-history traits are subject to the different selective pressures depending on the host’s developmental stage.

Previous studies have identified the adaptive significance of parasitizing early-stage hosts with delayed emergence (Slansky, 1986). First, young hosts are more abundant (and thus more available) than older hosts and thus are easier for wasps to locate (Price 1972). Second, young hosts are more susceptible to parasitism due to their immature defense mechanisms, such as thinner cuticles and weaker immune responses (Beckage and Riddiford 1978; Vinson and Iwantsch 1980b). Third, attacking young hosts can repel subsequent, heterospecific competitors attempting to attack the same hosts (e.g., in *Parasetigena silvestris* and *Blepharipa pratensis* parasitizing; Godwin and Odell 1984). Furthermore, delayed emergence may potentially increase their fecundity of the parasitoid, because it allows parasitoids to get more resource from their hosts (Harvey et al. 1994, 2004). On the other hand, delayed emergence is subject to a cost of increased mortality during the period for the host to grow (Price 1972; c.f., the “Slow-Growth High-Mortality hypothesis”; Clancy and Price, 1987; Benrey and Denno 1997). Godfray (1994) hypothesizes that the divergent selective pressures imposed by parasitizing early and late-stage hosts lead to the polarization implied by the idiobiont-koinobiont dichotomy.

Consistent evidence is found in species, e.g., *Hyposoter horticola*, with a minute ovipositor to lay eggs on the host (*Melitaea cinxia*) while pausing host development for approximately one year until the host becomes the seventh instar (Castelo et al., 2010). Thus, the adaptive significance of delayed emergence is qualitatively understood.

On the other hand, it remains unclear which factors such as host mortality, growth rates, and fecundity with host stage determine the directional selection for koinobiont versus idiobiont. Price (1972) constructed a framework in which the offspring number (*F*) is determined by the product of the number of offspring per female (*N*) and the survival probability of parasitoid larvae on host (*P*); namely, *F* = *N* ⋅ *P*. Price’s (1972) formula is a general but verbal model in the sense that ecological or physiological determinants of *N* and *P* are left unspecified, which limits the quantitative predictions of the direction of natural selection. Hackett-Jones et al. (2011) formulated the adaptive significance of delayed emergence. However, their model does not explicitly incorporate the stage-dependent traits of the host such as density ratio and parasitoid fecundities, the mortalities. These factors are critical for understanding the koinobiont strategy, since early-stage hosts often lack sufficient resources to support immediate emergence. Therefore, the key driving factor of evolution of koinobiont or idiobiont is yet to be better understood.

In this study, we use mathematical models to explore the adaptive evolution of delayed emergence. We focus on explaining the evolutionary dynamics of the dichotomy, assessing what conditions favor koinobiont versus idiobiont. To simplify the conceptualization of the results, we take operational definitions of idiobionts which attack late-stage hosts and consume immediately. We consider the evolutionary dynamics of two traits: (i) the parasitoids’ preference for hosts’ developmental stages, and (ii) the degree of delayed emergence for those parasitoids attacking young hosts. Using adaptive dynamics theory (Dieckmann and Law 1996), we determine the directions of natural selection on these traits and the stability of evolutionary consequences (i.e., the attainable strategies that no other rare mutant with small phenotypic differences can increase in frequency). As the host population is physiologically structured, we use Fisher’s reproductive values to interpret the optimal timing of emergence for parasitoids (Fisher 1930).

## Model

We first construct the dynamical equations of stage-structured host-parasitoid populations. We incorporate delayed emergence using a stage-structured model consisting of early-stage hosts, late-stage hosts, and adult hosts. Either stage of the stage hosts, but not adults, may be attacked by the parasitoids and subsequently transits to the parasitized stage (Fig 1 for a diagram depicting the patterns of life cycles). The variables and parameters are summarized in Table 1.

**Fig. 1.**
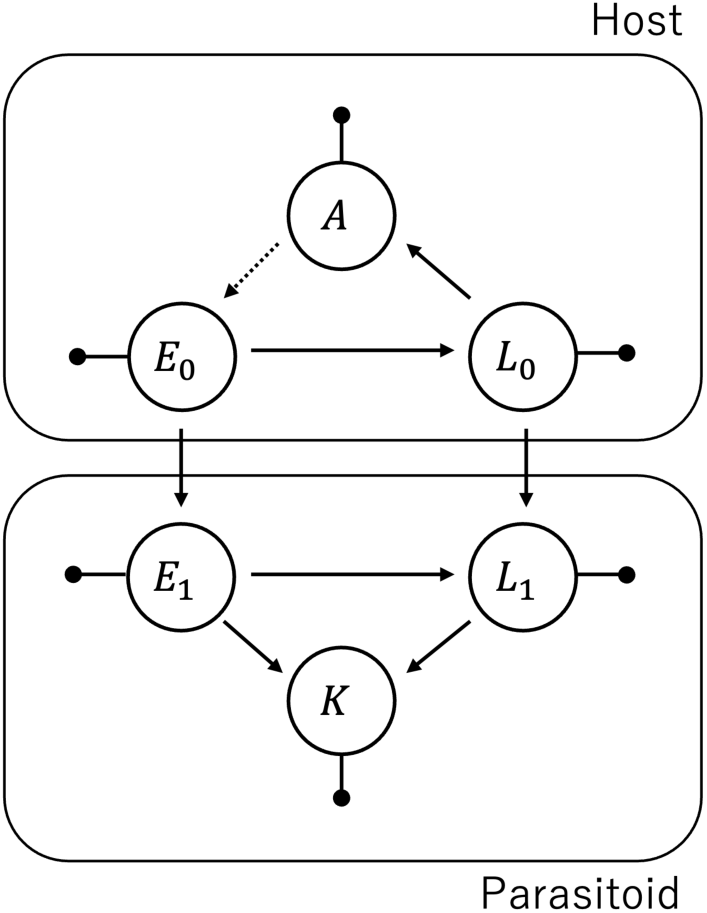
Diagram of host and parasitoids. Symbols: *E* for the density of early-stage hosts, *L* for the density of late-stage hosts, *A* for the density of adult hosts, and *K* for the density of parasitoids. We use subscript 0 for unparasitized and 1 for parasitized state, respectively. Arrows with black points represent death

**Table 1.**
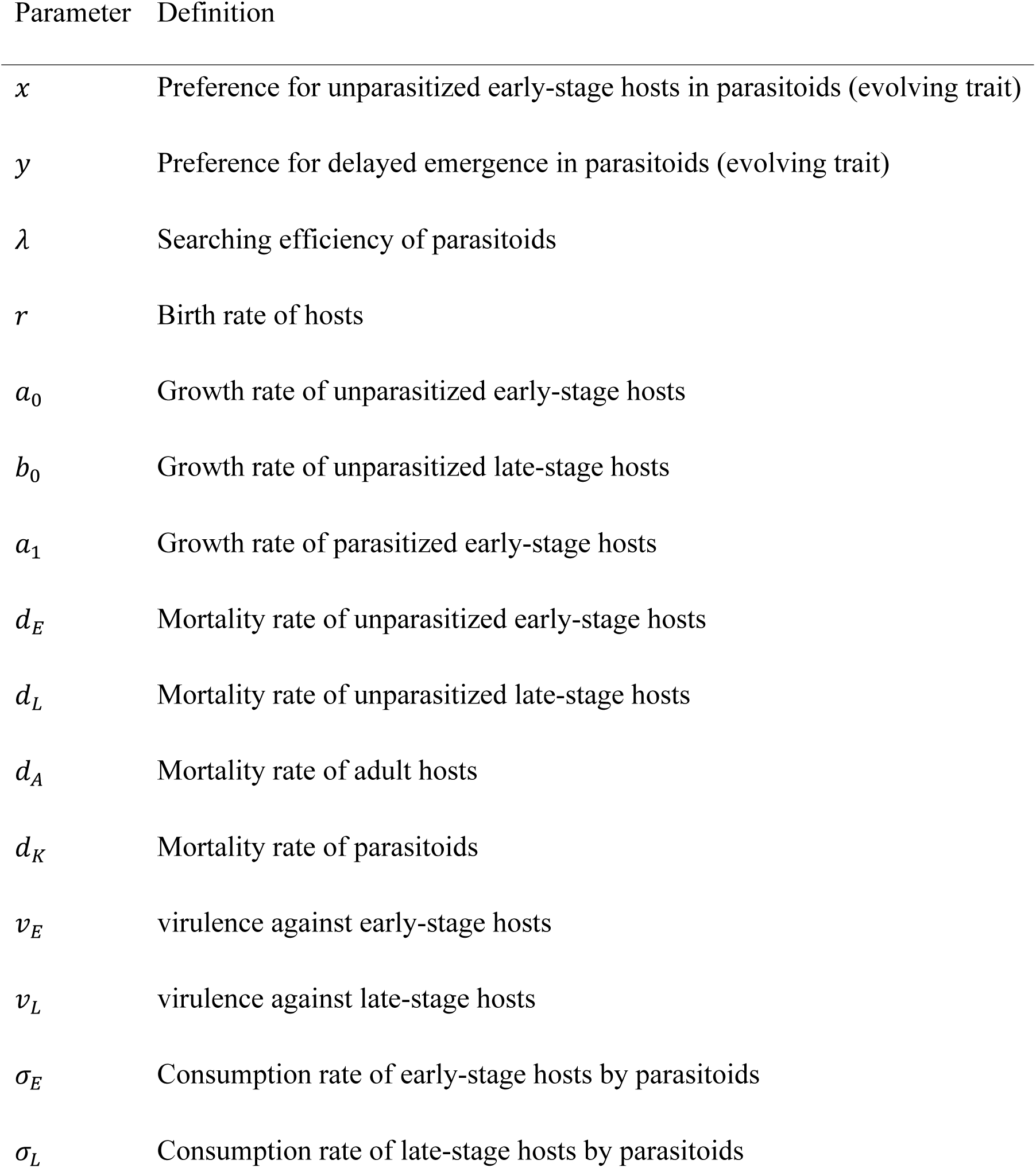
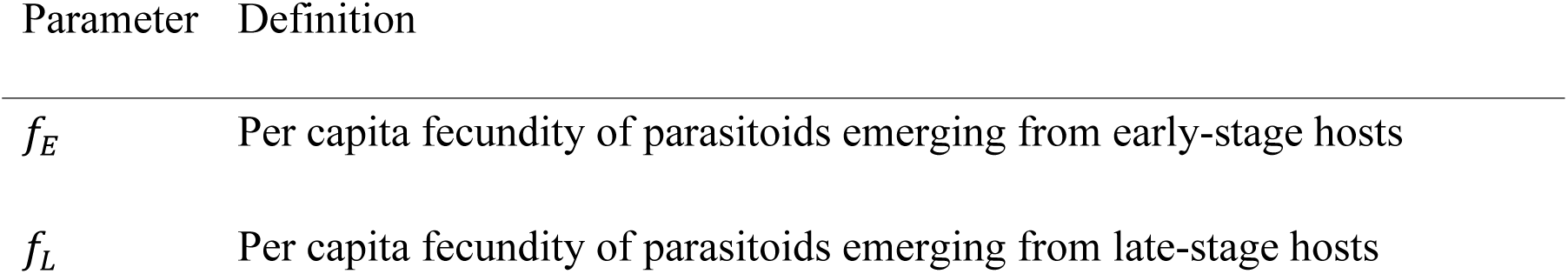
Definitions of the parameters.

| Parameter | Definition |
| --- | --- |
| $x$ | Preference for unparasitized early-stage hosts in parasitoids (evolving trait) |
| $y$ | Preference for delayed emergence in parasitoids (evolving trait) |
| $\lambda$ | Searching efficiency of parasitoids |
| $r$ | Birth rate of hosts |
| $a_0$ | Growth rate of unparasitized early-stage hosts |
| $b_0$ | Growth rate of unparasitized late-stage hosts |
| $a_1$ | Growth rate of parasitized early-stage hosts |
| $d_E$ | Mortality rate of unparasitized early-stage hosts |
| $d_L$ | Mortality rate of unparasitized late-stage hosts |
| $d_A$ | Mortality rate of adult hosts |
| $d_K$ | Mortality rate of parasitoids |
| $v_E$ | virulence against early-stage hosts |
| $v_L$ | virulence against late-stage hosts |
| $\sigma_E$ | Consumption rate of early-stage hosts by parasitoids |
| $\sigma_L$ | Consumption rate of late-stage hosts by parasitoids |

We consider two independent traits subject to natural selection. One is preference for early-stage hosts, *x*, and the other is probability of delayed emergence, *y*, with 0 ≤ *x* ≤ 1 and 0 ≤ *y* ≤ 1. For example, the case with *x* = 1 (attacking early-stage hosts only) and *y* = 0 (emerging only from early-stage hosts) corresponds to the life cycle graph in Fig 2A, which thus omits the lifecycle pathway between early-stage and late-stage hosts. We do not consider the evolution of host traits.

**Fig. 2.**
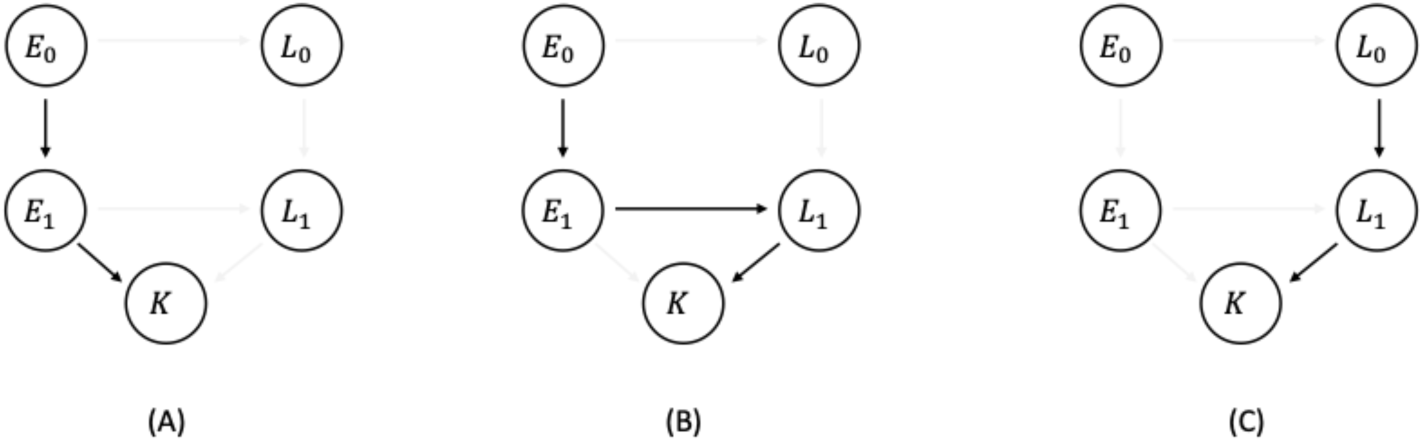
Three pathways of reproductive success: (A) attacking the early-stage hosts and immediately preying on them; (B) attacking the early-stage hosts and waiting until they grow to late-stage hosts, and then preying on them; (C) attacking the late-stage host and immediately preying on them

We use ordinary differential equations to describe the population dynamics of hosts and parasitoids:

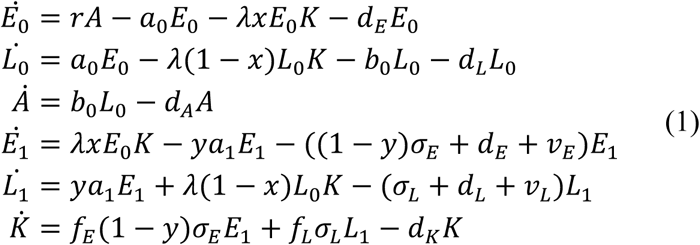

(with dot representing the time derivative). Here, from top to bottom, the ordinary differential equations represent the dynamics of non-parasitized early-stage host (*E*_0_), non-parasitized late-stage hosts (*L*_0_), adult hosts (*A*), parasitized early-stage host (*E*_1_), parasitized late-stage host (*L*_1_), and adult parasitoids (emerging from either early or late hosts; *K*). The key ingredients are the transitions in states due to (i) parasitism (e.g., from *E*_0_ to *E*_1_), and (ii) ontogeny (e.g., from *E* to *L*) (Fig 1). Surviving late-stage hosts may develop into adults (with density *A*).

The first three lines in Equation (1) describe the population dynamics of non-parasitized hosts. With non-parasitized early-stage hosts parasitized at rate *λxK* with searching efficiency of parasitoids *λ* and the probability of preferentially parasitizing early-stage hosts *x*, the non-parasitized early-stage hosts (with density *E*_0_) are produced by adults at rate *r*, develop into late-stage hosts at rate *a*_0_ and die at rate *d*_*E*_; they are parasitized by wild-type parasitoids at rate *λx*. Non-parasitized late-stage hosts (with density *L*_0_) develop from non-parasitized early-stage hosts but may be parasitized at rate *λ*(1 − *x*)*K*, develop into adults at rate *b*_0_, or die at rate *d*_*L*_.

The middle two lines in Equation (1), on the other hand, describe the population dynamics of parasitized hosts. Parasitized early-stage hosts (with density *E*_1_), after transiting from the parasitism of early-stage hosts (*E*_0_), develop into late-stage hosts, or otherwise die due either to (i) parasitoid emergence, at rate (1 − *y*)𝜎_*L*_ (ii) natural mortality at rate *d*_*E*_, or (iii) virulence at rate 𝑣_*E*_ (increased mortality due to parasitism). Parasitized late-stage hosts (with density *L*_1_) develop from parasitized early-stage hosts due to ontogeny or transit from non-parasitized late-stage hosts due to parasitism but die at rate 𝜎_*L*_ due to emergence of the parasitoid, or at rate *d*_*E*_ + 𝛾_*E*_ due to natural mortality or virulence. We assume that the parasitized hosts are incapable of producing offspring.

The last line in Equation (1) describes the population dynamics of adult parasitoids. Adult parasitoids emerge from either parasitized early-stage hosts (at rate 𝑓_*E*_) or late-stage hosts (at rate 𝑓_*L*_), where 𝑓_*E*_ or 𝑓_*L*_ represents the fecundity of adult parasitoids (per capita) emerging from early larvae or late larvae, respectively.

To investigate the long-term evolutionary dynamics of the preference for host stages in parasitoids and the preference for delayed emergence in parasitoids, we apply the adaptive dynamics theory (Dieckmann and Law 1996). We first suppose that the resident population dynamics, Equation (1), have reached an endemic equilibrium, and introduce a rare mutant of the parasitoid attempting to invade the resident equilibrium. The resident population has a unique equilibrium (Appendix D). To determine whether the mutant parasitoid can invade the resident population, we derive the invasion fitness proxy 𝑅_m_ by using the next-generation theorem (Van den Driessche and Watmough 2002; Hurford et al. 2010):

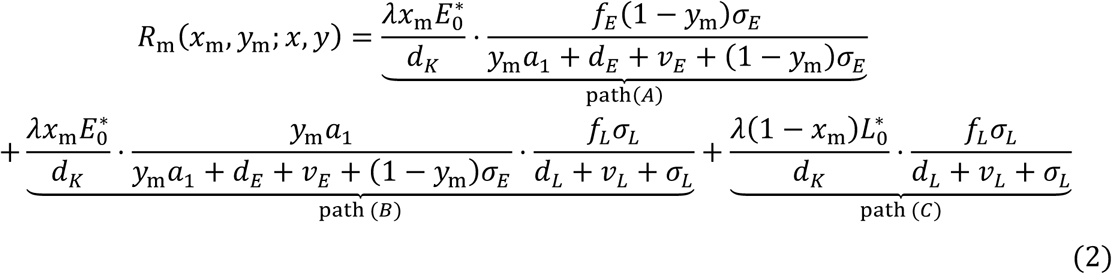

where *E*^∗^_0_ and *L*^∗^_0_ represent the equilibrium density of unparasitized early-stage host and unparasitized late-stage host, respectively. In Equation (2), we partition the reproductive success of the mutant individual into multiple pathways (Fig 2). The first, path (A) term represents the reproductive success by attacking and consuming early-stage hosts (Fig 2A). The path (B) term represents the reproductive success by attacking on early-stage hosts and consuming late-stage hosts (Fig 2B). The path (C) term represents the reproductive success by attacking and consuming late-stage hosts (Fig 2C).

The rare mutant can invade the resident population when 𝑅_m_ > 1. By using this criterion, we examine a candidate evolutionary consequence, its evolutionary stability (i.e., whether the rare mutant can invade a population dominated by the wild type; Maynard Smith 1982), and also convergence stability (i.e., evolutionary dynamics convergence of the traits; Eshel et al. 1983).

Although the results in the main text all assume the exponential growth of the parasitoid-free host population (as this assumption immensely simplify the analyses), we also analyze the case where host density follows logistic growth (i.e., density-dependent regulation of host fecundity). To check the robustness of the results in the main text, we conduct numerical, deterministic simulations (Appendix E).

## Results

### Evolution of delayed emergence

First, we present one of the key results that the delayed emergence trait *y* is polarized; i.e., natural selection pushes the population towards either *y* = 0 (immediate emergence) or *y* = 1 (complete delayed emergence). We assess the directional selection on the preference for delayed emergence *y*, by using the selection gradient (Appendix B) given by:

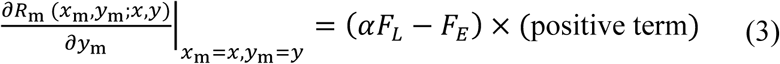

which is positive (or negative) when natural selection favours delayed (or early) emergence (respectively), depending on the sign of the first term, (𝛼*F*_*L*_ − *F*_*E*_). In Equation (3), 𝛼 = *a*_1_/(*a*_1_ + *d*_*E*_ + 𝑣_*E*_) is the probability of successful development from early to late-stage for parasitized host, *F*_*L*_ = 𝑓_*L*_𝜎_*L*_/(*d*_*L*_ + 𝑣_*L*_ + 𝜎_*L*_) or *F*_*E*_ = 𝑓_*E*_𝜎_*E*_/(*d*_*E*_ + 𝑣_*E*_ + 𝜎_*E*_) is the product of: (1) the probability of consumption of the host and (2) the fecundity (i.e., the expected conversion rate of parasitoids preying on early or late-stage hosts, respectively). Although the selection gradient per se depends on both of the evolutionary traits *x* and *y*, the sign is determined only by non-evolving parameters such as 𝛼 and *F*_*L*_. Thus, the preference evolves monotonically to either 0 or 1 (i.e. immediate or delayed emergence) depending on the sign of 𝛼*F*_*L*_ − *F*_*E*_. We also find that the evolutionary outcome is always evolutionarily and convergence stable (Appendix C), and therefore Equation (3) predicts that the long-term evolutionary outcome is either immediate emergence or completely delayed emergence (Otto and Day 2007).

We can interpret the monotonicity result of Equation (3) using the Fisher’s (1930) reproductive values. Equation (3) represents the relationship between the reproductive value of early-stage hosts when the parasitoids consume immediately versus when they wait for the host to develop into late-stage hosts before feeding. Here, the parameters 𝛼*F*_*L*_ − *F*_*E*_ quantify the balance between reproductive gains of parasitoids from these two strategies: 𝛼*F*_*L*_ represents the expected number of adult parasitoids when the host survives the initial attack, grows to the late-stage stage, and is consumed at that stage (pathway B in Fig 2), whereas *F*_*E*_ measures the number of adult parasitoids that can be produced when an early-stage host is consumed immediately (pathway A in Fig 2). The preference for delayed emergence is likely to be favoured when the product of (1) the growth probability of the parasitized early-stage hosts and (2) the average conversion rate when parasitoids consume the late-stage host, is higher than (3) the conversion rate of the early-stage host. These conditions mean that delayed emergence is favoured when late-stage hosts are overall more valuable as resource for parasitoids than early-stage hosts. Taken together, our models predict that the delayed emergence trait is polarized, leading to a clear division of parasitoids into koinobionts and idiobionts depending on these conditions.

**Fig. 3.**
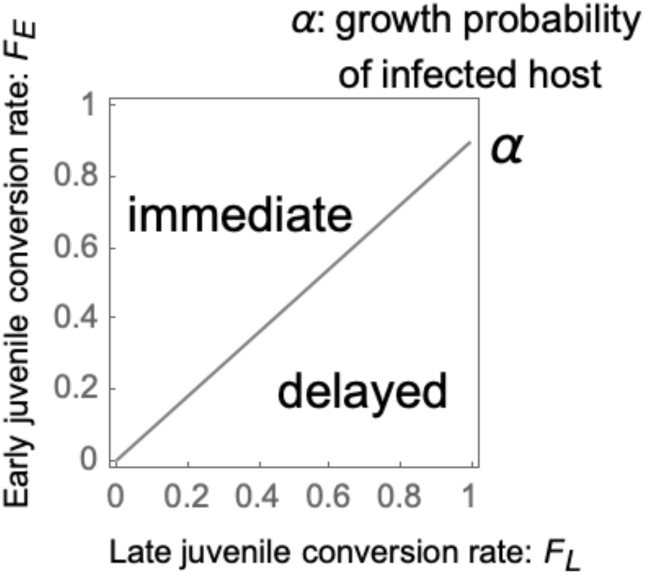
The evolutionary consequences of the preference for delayed emergence (𝛼 = *a*_1_/(*a*_1_ + *d*_*E*_ + 𝑣_*E*_): the growth probability of the parasitized early-stage host, *F*_*E*_ = 𝑓_*E*_𝜎_*E*_/(*d*_*E*_ + 𝑣_*E*_ + 𝜎_*E*_): the average fecundity rate of parasitoids preying on the early-stage host and *F*_*L*_ = 𝑓_*L*_𝜎_*L*_/(*d*_*L*_ + 𝑣_*L*_ + 𝜎_*L*_): that on the late-stage host)

### Patterns of life cycles

Next we examine the evolution of the preference to attack early-stage hosts (*x*). The selection gradient of the preference for host stages *x* is given by:

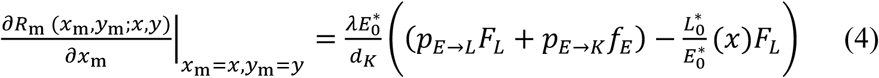

(Appendix B), where the compound parameter 𝑝_*E*→*L*_ = *ya*_1_/(*ya*_1_ + *d*_*E*_ + 𝑣_*E*_ + (1 − *y*)𝜎_*E*_) represents the survival probability (transition) from parasitized early-stage hosts to late ones and 𝑝_*E*→*K*_ = (1 − *y*)𝜎_*E*_/(*ya*_1_ + *d*_*E*_ + 𝑣_*E*_ + (1 − *y*)𝜎_*E*_) represents the probability of parasitoid emergence among dead hosts The sign of Equation (4) depends on both *x* and *y*; note that according to the previous section, *y* evolves to either 0 or 1 regardless of the evolution of *x*.

Now we substitute *y* = 0 or 1 considering the result in the previous section. The selection gradients on the preference for early-stage hosts for both cases are given by:

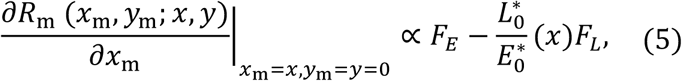

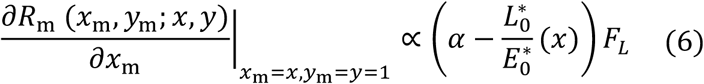

where *F*_*E*_ = 𝑓_*E*_𝜎_*E*_/(*d*_*E*_ + 𝑣_*E*_ + 𝜎_*E*_) is the product of (1) the probability of consumption of early-stage hosts and (2) the fecundity of them (i.e., the expected fecundity of parasitoids preying on early-stage hosts) and *F*_*L*_ = 𝑓_*L*_𝜎_*L*_/(*d*_*L*_ + 𝑣_*L*_ + 𝜎_*L*_) is that on late-stage host, 𝛼 = *a*_1_/(*a*_1_ + *d*_*E*_ + 𝑣_*E*_) is the probability of successful development of parasitized early-stage host to late-stage host. We also find that the evolutionary outcome is always evolutionarily and convergence stable (Appendix C), and therefore Equation (5) and (6) predicts the long-term evolutionary outcome (Appendix E). A unique candidate evolutionary consequence *x*, obtained analytically from Equations (5) and (6), is defined as the singular point where the selection gradient becomes zero. When the candidate evolutionary consequence falls outside the interval between 0 and 1, we carry out case-analyses to locate the interior singular point (Appendix B.3). The evolutionary consequence in each case is shown in Fig 4. This result suggests that delayed emergence trait can generate diverse life cycles of parasitoids, including idiobiont (Fig 4 (D-v)) koinobiont (Fig 4 (D-ii) and (D-iv)) (see below). Figs 4 (D-ⅰ) and (D-ⅲ) exhibit strategies which immediate exploit early-stage hosts. These patterns are biologically implausible since early-stage hosts provide insufficient resources for parasitoid development.

**Fig. 4.**
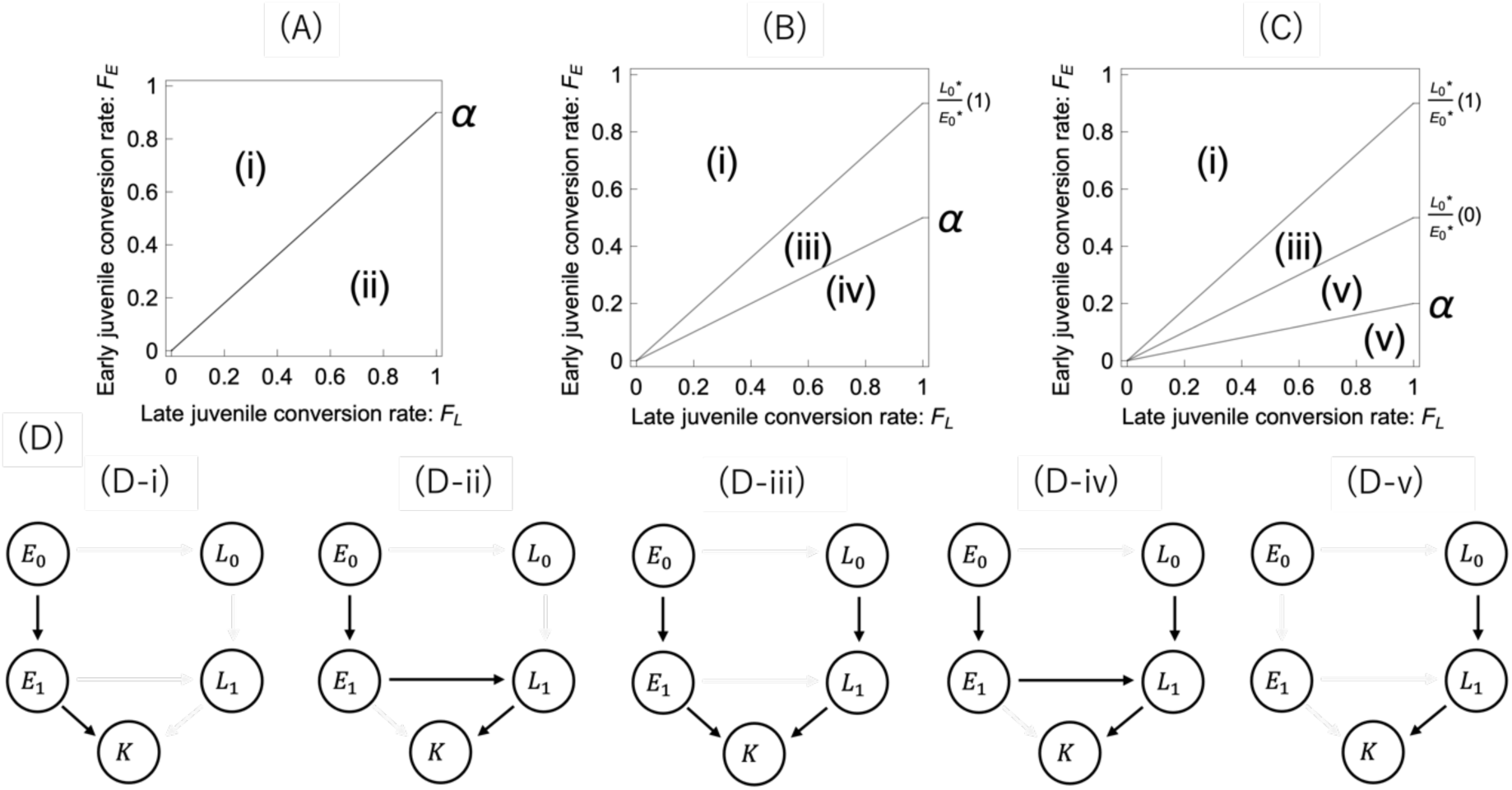
The patterns of delayed emergence resulting from our model. (A) Only early hosts are attacked (the singular point is 0) when the growth probability of the parasitized early-stage host 𝛼 is larger than the maximum of the density ratio of the unparasitized late-stage host to the early-stage hosts *L*^∗^_0_ /*E*^∗^_0_. (B) Both early and late hosts are attacked (the singular point is between 0 and 1) when the growth probability of the parasitized early-stage host 𝛼 is between the minimum and maximum of *L*^∗^_0_ /*E*^∗^_0_). (C) Only late hosts are attacked (the singular point is 1) when 𝛼 is smaller the minimum of *L*^∗^_0_ /*E*^∗^_0_). (D) The five panels show distinct evolutionary outcomes in parasitoid life cycles resulting from variation in emergence timing and host stage exploitation, referred to as (ⅰ)-(ⅴ) in the graphs. *E* or *L* represents early-stage hosts or late-stage hosts (respectively). (ⅰ) only early hosts are attacked and they are consumed immediately. (ⅱ) only early hosts are attacked and they are not consumed immediately. (ⅲ) both early and late hosts are attacked and they are consumed immediately. (ⅳ) both early and late hosts are attacked and late ones are consumed. (ⅴ) late hosts are attacked and they are consumed immediately

### Evolution of idiobiont and koinobiont

Following Godfray (1994), we define idiobiont as the parasitoids attacking unparasitized late-stage hosts and prey on the host immediately. Similarly, we define koinobiont as the parasitoids that attack an unparasitized early-stage host and prey on the host after the host grows up to the late-stage stage (Godfray, 1994). Therefore, Fig 4 (D-ii) represents koinobiont and Fig 4 (D-v) represents idiobiont. The evolution of koiobionts is more likely when the survival probability of parasitized early-stage hosts to late ones is high, and the relative density of late-stage hosts to early-stage hosts is small. Fig (D-iv) provides a clear illustration of the strategy depending on host stage at parasitism in koinobionts. While koinobionts can attack hosts at various developmental stages, they differentiate their approach—either allowing further host growth or initiating immediate consumption—depending on the specific stage encountered. The evolution of idiobionts is more likely when the survival probability of parasitized early-stage hosts to late ones is low, and the relative density of late-stage hosts to early-stage hosts is large.

Finally, we check the robustness of the results based on the exponential growth of hosts in the absence of parasitoid. Numerical simulations (assuming the host population exhibits logistic growth due to density-dependent regulation) confirmed the robustness of our results against the logistic growth case (Appendix E).

## Discussion

In this study, we investigated the effect of stage-structure and host ecology on the evolution of life-history traits in parasitoids. Our primary achievement is the formulation of the selection pressure on the delayed emergence using the concept of reproductive value (Fisher 1930). Specifically, if the host’s developmental growth does not increase the expected conversion efficiency (a measure of nutritional quality), where *F*_*L*_ ≤ *F*_*E*_, the parasitoids are then selected to emerge from the host immediately (Fig 3). Otherwise, either early or late emergence is selected for (Fig 3). The specific outcome of this selection is determined by comparing the reproductive value of the host for parasitoids emerging early (*F*_*E*_) with that for those waiting for host growth (𝛼*F*_*L*_) (Fig 3). Moreover, despite its fundamental structure, our model can predict diverse evolutionary outcomes of parasitoid life cycles (Fig 4). To incorporate delayed emergence into the model, we considered two developmental stages of hosts available for parasitism, allowing parasitized early-stage hosts to continue developing into later stages. As a result, the host utilization strategies of parasitoids exhibit five distinct patterns (Fig 4D). This result indicates that the evolution of emergence timing leads to the observed diversity of parasitoid life cycles, including both idiobiont- and koinobiont-strategies. Notably, our model can account for the strategy observed in many koinobionts, which utilize both early- and late-stage hosts (Fig 4 (D-iv)). When parasitoids attack early-stage hosts, the model predicts that they delay emergence to maximize resource acquisition from host growth. Conversely, when parasitoids attack late-stage hosts, our model predicts immediate consumption. This result aligns with the patterns in which the duration of delayed emergence is shortened when the host already has high resource value. This strategy involves a dynamic preference between host stages, driven by their relative abundance (density ratio). Our model explores how the trade-offs between host stages shift depending on the density ratio. This finding provides a theoretical explanation for why koinobionts do not always delay their emergence but instead finetune the timing based on the host’s value at the time of parasitism.

Our predictions are overall consistent with a previous theoretical study on stage-structured host populations. Iritani et al. (2019) considered stage-structured disease dynamics and showed that the evolutionary consequences of virulence toward larval hosts depend on the interaction between host maturation rate and the infection pathway. Specifically, virulence toward larval hosts evolves to be higher when either (i) the adult stage is relatively long and the transmission across host stages mostly occurs, or (ii) the larval stage is relatively long and the transmission across the same stage mostly occurs (Iritani et al. 2019). These findings align with our results: koinobiont strategies evolve when early-stage hosts are abundant and maturation under infection is likely, whereas the idiobiont strategy evolves when late-stage hosts are abundant and maturation is unlikely. Hackett-Jones et al. (2011) investigated the evolution of delayed emergence in parasitoids using a discrete-time model, showing that delayed emergence is under balancing selection between two forces. First, delayed emergence allows parasitoids to attain a larger adult body size, which is favored by the increased searching efficiency associated with larger size. Second, delayed emergence is costly because of the host mortality during the period of delayed emergence. In this context, our models are a simpler generalization of their model for two reasons. Compared to Hackett-Jones et al.’s (2011) model assuming the host population is homogeneous (thus with no structure), we explicitly consider host stage-structure. Also, the authors’ analytical formula involves host density in a complex mathematical form, making biological interpretation difficult. Even more crucially, their model does not account for two key biological facts: (1) young hosts are more abundant than older hosts (Price, 1972), and (2) young hosts are easier to parasitize due to weaker resistance, such as lower physical defense or immunity (Beckage and Riddiford 1978; Vinson and Iwantsch 1980b). Our model incorporates these essential factors to derive a biologically interpretable formula of selection pressure for delayed emergence, revealing the optimal strategy is determined by the relative reproductive value of host stages depending on the balance between host growth, mortality, and nutritional quality.

Our model predicts that host density (density ratio) has strong impact on the evolution of delayed emergence. In environments where early-stage hosts are relatively abundant compared to late-stage hosts, natural selection favours delayed emergence which exploits the high encounter rate of early-stage hosts and waits for growth (Slansky 1986). In nature, early-stage hosts are more numerous than late-stage hosts. Therefore, even if there is a mortality cost while waiting for growth within the host, the benefits provided by the high encounter rate can outweigh this cost. Conversely, idiobionts tend to evolve in environments where the natural mortality of the host is low and late-stage hosts are nearly as abundant as early-stage hosts. This allows them to efficiently avoid the cumulative mortality associated with the period required for early-stage hosts to mature (Price 1972).

Host density best illuminates ecological characteristics of the host (host ecology). We discuss this point focusing on host natural mortality caused by scavengers and other predators. Idiobionts tend to attack concealed hosts, such as those within plant tissues or cocoons (Price 1972; Gauld, 1988; Quicke, 2014). This preference arises because permanent paralysis by idiobiont renders the host entirely defenseless. In an exposed environment, such an incapacitated host and the parasitoid larva feeding on it would be highly vulnerable to natural mortality from scavengers and predators (Gauld, 1988). Idiobionts target hosts in concealed niches where this natural mortality 𝑣_*L*_ is sufficiently small (Gauld, 1988). In addition, late-stage hosts are likely to be relatively abundant in these protected environments because natural mortality due to predation throughout the developmental period *d*_*E*_ is relatively low (Price, 1972). These aspects of host ecology support the association between immediate emergence and the parasitization of late-stage hosts, under the selection pressures on parasitoids defined by Equation (3) for delayed emergence and Equation (4) for host stage, respectively. In contrast, koinobionts often attack exposed hosts that feed on surfaces such as leaves (Price 1972; Gauld, 1988; Quicke, 2014). Paralyzing hosts in such a location would lead to an increase in natural mortality, as the host and the parasitoid larva feeding on it would be easily discovered and preyed upon (Gauld, 1988). Therefore, maintaining host mobility is an essential benefit to ensure the parasitoid’s survival (Gauld, 1988). Koinobionts maintain host mobility so that this natural mortality 𝑣_*E*_ remains sufficiently small even in exposed niches (Gauld, 1988). Furthermore, in these exposed environments, late-stage hosts are rarely abundant because high natural mortality *d*_*E*_ throughout development limits the number of individuals reaching maturity (Price 1972). These aspects of host ecology support the association between delayed emergence and the parasitization of early-stage hosts in koinobionts, under the selection pressures defined by Equation (3) for delayed emergence and Equation (4) for host stage, respectively. These formulations explain how host ecology, specifically stage-specific mortality, leads to the polarization between idiobiont and koinobiont.

The present results predict that the increase in fecundity as the host matures favours delayed emergence. In fact, the positive correlation between host size and fitness-related traits such as adult size, egg count, and longevity is particularly pronounced, because they halt host growth (Salt 1941, Arthur and Wylie 1959, Vinson 1972, Sandlan 1979, Strand et al. 1988, King 1989). However, in koinobionts, a larger host size does not necessarily translate into a positive impact on fitness-related traits (Harvey, 2016). The host is not merely food; the parasitoid must highly synchronize its growth with the host’s physiological and developmental stage (Harvey, 2016). For example, *Apanteles carpatus* attacks hosts (*Tineola bisselliella*) which are as small as 3% (30–40 mg) of its own body weight (1–1.5 mg). While it takes about 30 days for the parasitoid to develop from a fifth-instar host to adult stage, it takes up to 87 days for the parasitoid to develop from a first– third-instar host to adult stage (Harvey, 2000). On the other hand, there are negative factors such as the failure to pupate when they cannot fully consume excessively large hosts (Harvey 1996) and the host’s strong immune response in older hosts (Harvey et al. 2004). That is, host’s value to koinobionts must be understood not as being directly proportional to host size at the time of oviposition (Sequeira and Mackauer 1992a,b; Harvey 2012). Therefore, the idiobiont-koinobiont dichotomy should be considered as a continuum because koinobionts can flexibly adjust the degree of delayed emergence depending on the host’s developmental stage at the time of parasitism. Thus, future theoretical studies would benefit from taking these processes into account to explain the life-history variations in parasitoid wasps.

We made some simplifying assumptions. For instance, the possibility of multiple parasitisms is omitted from the model, despite its occurrence in nature (Price, 1972; Harvey et al., 2013; Ode et al., 2022). Multiple parasitisms are associated with competition, which drives temporal niche differentiation in host developmental stages (Price, 1972). This niche differentiation promotes the evolution of delayed emergence (Price, 1972). Future theoretical studies may incorporate the effects of such competition, which may provide testable predictions for the evolution of life-history variations in parasitoids. Moreover, hosts can also evolve defense traits such as resistance, potentially leading to coevolution. Age structure of hosts may further shape these dynamics, as shown in a model of tolerance– virulence coevolution (Buckingham and Ashby 2024). Future extensions of the present model may incorporate host evolution to develop a useful theoretical framework for understanding how host–parasitoid coevolution drives variation in emergence timing and associated life-history traits.

In summary, this study rigorously formulated the selection pressure on delayed emergence in parasitoids using the concept of reproductive value, providing a theoretical foundation to make testable predictions for existing empirical findings. The primary contribution of this model lies in providing measurable quantitative indices for the idiobiont-koinobiont dichotomy. Future studies may extend our model to incorporate the adjustment of delayed emergence depending on the host’s developmental stage in koinobionts to understand the multi-layered dichotomy.

## Appendix

### A. Next generation matrix and invasion fitness proxy

To study life-history evolution, we assess the invasion possibility of a rare mutant into the resident population assuming that the ecological dynamics has reached an equilibrium. The population dynamics of the rare mutant is given by:

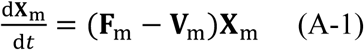

where

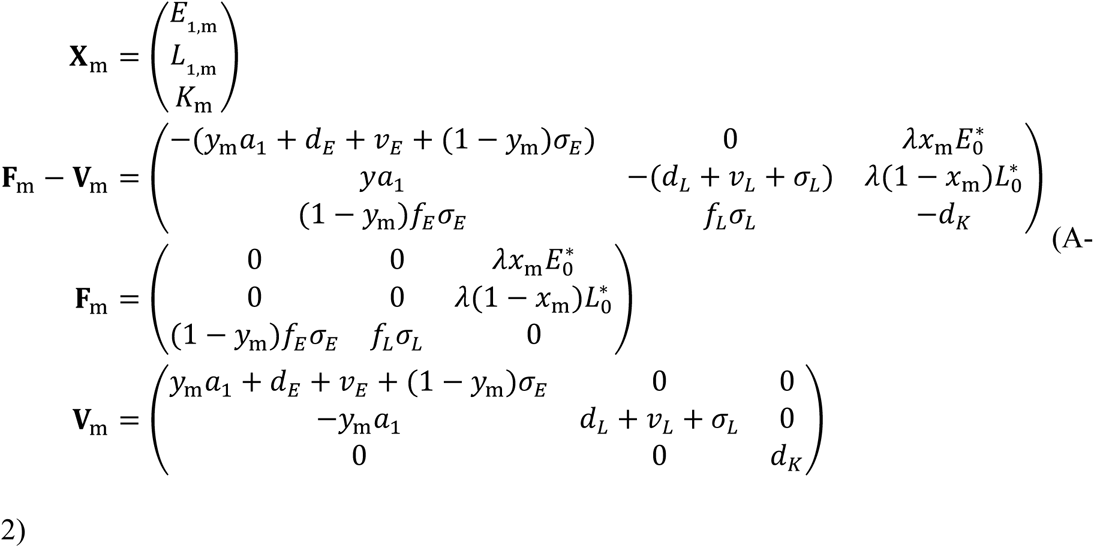

The Next-Generation Theorem (Van den Driessche and Watmough 2002) states that if the dominant eigenvalue of the next-generation matrix 𝐆_m_ = 𝐅_m_ ⋅ 𝐕_m_^−𝟏^ exceeds 1, then the mutant can invade and drive the resident individuals extinct. The squared, dominant eigenvalue 𝜌(𝐆_m_) is given by:

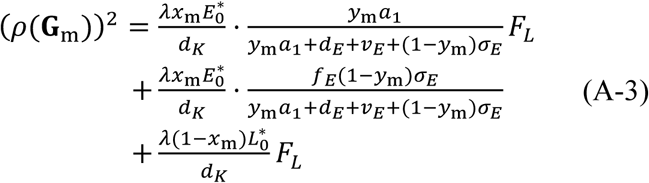

where

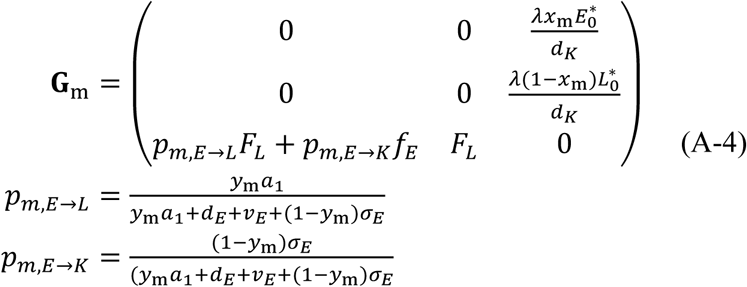

where 𝑝_𝑚,*E*→*L*_ represents the probability that mutant parasitoids’ larvae can transit from parasitized early-stage hosts to late ones and 𝑝_*E*→*K*_ represents the probability that mutant parasitoids’ larvae consume the host. Since 𝜌^2^ ≷ 1 is equivalent to 𝜌 ≷ 1, we use 𝜌^2^ for the invasion proxy.

### B. Selection gradient

In this section, we derive the stable class distribution, reproductive values and selection gradient using the next-generation matrix.

#### B-1. Stable class distribution

The right eigenvector associated with the dominant eigenvalue of the next-generation matrix gives the stable class distribution (Taylor 1990; Otto and Day 2007), which represents the asymptotic distribution of mutant alleles. It is given by:

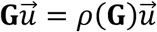

where

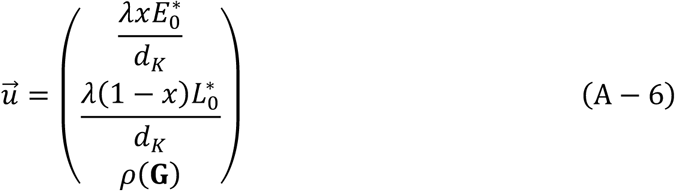

represents the stable distribution of the mutant allele.

#### B-2. Reproductive value of each class

The left eigenvector associated with the dominant eigenvalue of the next-generation matrix gives the reproductive values of each of these classes (Fisher 1930; Otto and Day 2007). The reproductive values represent the contributions of individuals in each class to future reproduction after sufficiently long generations, given by:

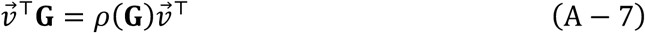

where

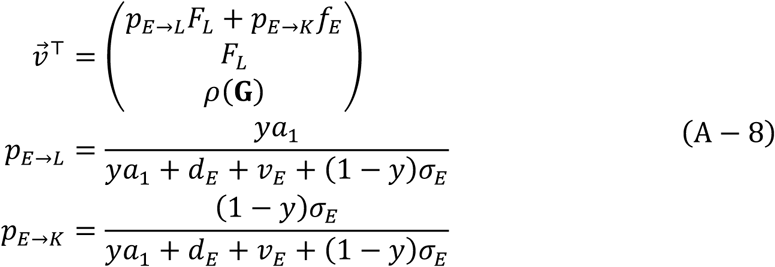

𝑝_*E*→*L*_ represents the probability that parasitoids’ larvae can transit from parasitized early-stage hosts to late ones and 𝑝_*E*→*K*_ represents the probability that mutant parasitoids’ larvae consume the host.

#### B-3. Selection gradient of *x*

From the calculations above, we have 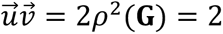. Let 𝑐_𝑖_ = 𝑢_𝑖_𝑣_𝑖_ be class reproductive value of class 𝑖; the selection gradient of *x* is given by:

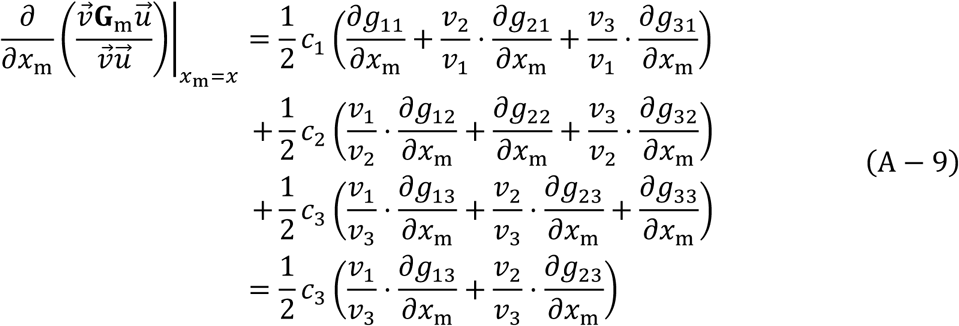

where

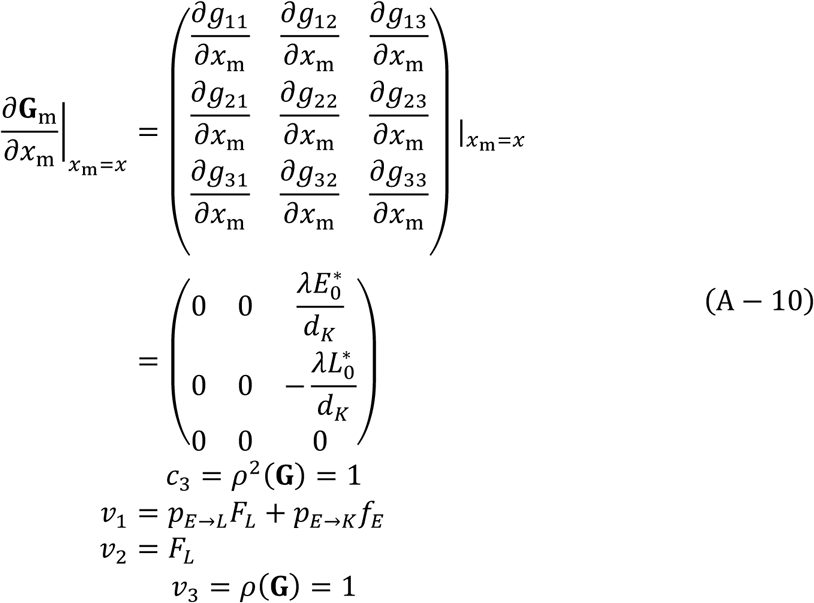

#### B-4. Evolution of preference for host stages when immediate emergence occurs

The selection gradient on the preference for early-stage hosts is

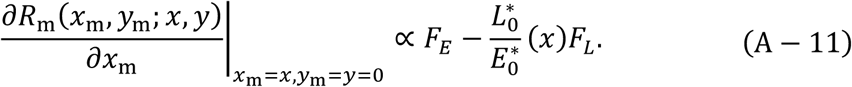

where *F*_*E*_ = 𝑓_*E*_𝜎_*E*_/(*d*_*E*_ + 𝑣_*E*_ + 𝜎_*E*_) is the expected fecundity of parasitoids preying on early-stage host and *F*_*L*_ = 𝑓_*L*_𝜎_*L*_/(*d*_*L*_ + 𝑣_*L*_ + 𝜎_*L*_) is that on late-stage host. A unique candidate evolutionary consequence *x*^∗^can be obtained analytically. The point is always convergence stable because *L*^∗^_0_ /*E*^∗^_0_ is monotonically increasing function of *x*^∗^ (Appendix D). Evolutionary consequences of the preference for host stages are always continuously stable in the case that parasitoids consume hosts immediately after they attack the hosts.

According to result section, we obtain the evolutionary consequences in three cases (Fig 5). Equation Equation (A-11) represents the relationship between the class reproductive value of unparasitized early-stage hosts and that of late ones when the parasitoids prey immediately. We can interpret Equation Equation (A-11) as the evolution of delayed emergence such that the hosts’ class reproductive value maximizes. When parasitoids consume hosts immediately after they attack hosts, the preference for early-stage hosts is likely to evolve if the average conversion rate and the density of late-stage hosts are higher than that of early-stage hosts. In addition, the lower the growth probability of parasitized early-stage hosts, the greater the diversity of host stages can evolve.

**Fig. 5.**
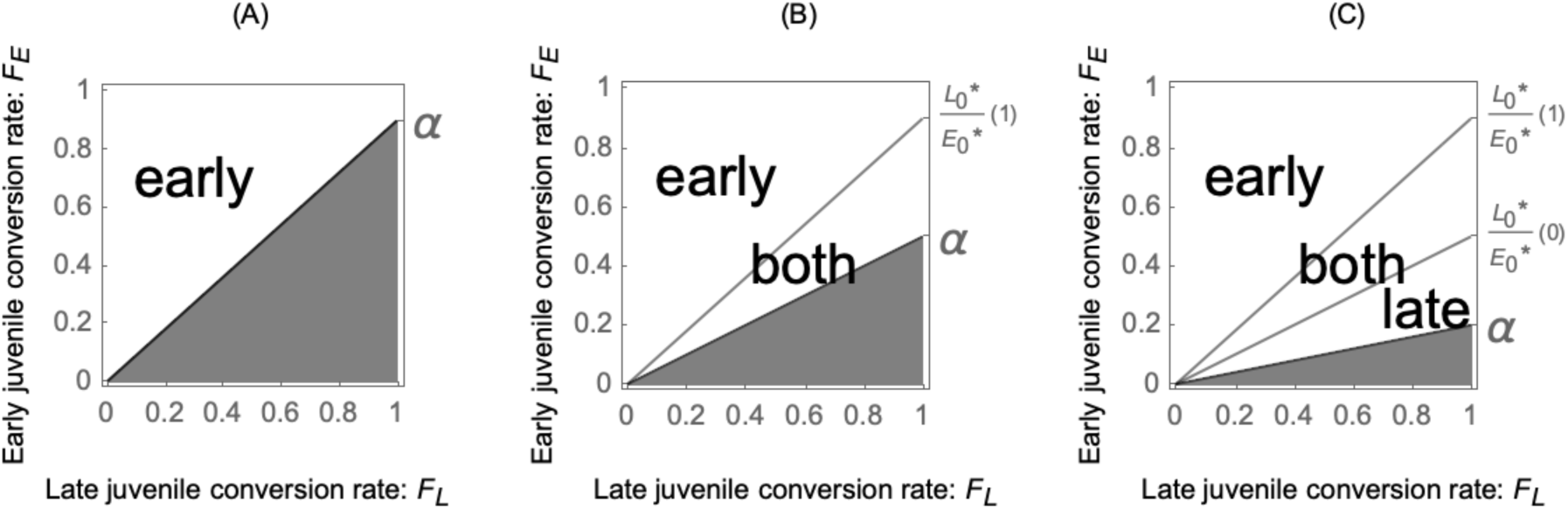
Evolutionary consequences of the preference for host stages when parasitoids emerge immediately. Immediately emerge ( *y* = 0) is convergence stable in the white area, but not in the gray area. Parasitoids attack only early-stage hosts in the “early” area, attack both of early and late-stage hosts in the “both” area, and attack late-stage hosts in the “late” area. (a) The singular point is 0 (the growth probability of the parasitized early-stage host 𝛼 is larger than the maximum of the density ratio of the unparasitized late-stage host to the early one *L*^∗^_0_ /*E*^∗^_0_). (b) The singular point is between 0 and 1 (𝛼 is between the minimum and maximum of *L*^∗^_0_ /*E*^∗^_0_). (c) The singular point is 1 (𝛼 is smaller the minimum of *L*^∗^_0_/*E*^∗^_0_)

#### B-5. Evolution of preference for host stages when delayed emergence occurs

The selection gradient on the preference for early-stage hosts is

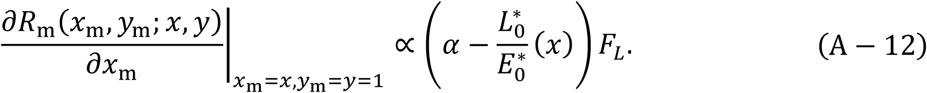

where 𝛼 = *a*_1_/(*a*_1_ + *d*_*E*_ + 𝑣_*E*_) is the growth probability of parasitized early-stage host, *F*_*L*_ = 𝑓_*L*_𝜎_*L*_/(*d*_*L*_ + 𝑣_*L*_ + 𝜎_*L*_) is the expected fecundity rate of parasitoids preying on late-stage host. A unique candidate evolutionary consequence *x*^∗^can be obtained analytically. The point is always convergence stable because *L*^∗^_0_ /*E*^∗^_0_ is monotonically increasing function of *x*^∗^ (Appendix D). Evolutionary consequences of the preference for host stages are always continuously stable in the case that parasitoids do not consume hosts immediately after they attack the hosts. According to result section, we obtain the evolutionary consequences in three cases (Fig 6). Equation Equation (A-12) represents the relationship between the class reproductive value of unparasitized early-stage hosts and that of late ones when parasitoids wait for their maturation. We can interpret Equation Equation (A-12) as the evolution of delayed emergence such that the hosts’ class reproductive value maximizes. Therefore, in the case that parasitoids do not consume hosts immediately after they attack the hosts, the preference for the unparasitized early-stage host is likely to evolve under two conditions: (1) the growth probability of the parasitized early-stage host is high, and (2) the density of the late-stage host is lower than that of early-stage hosts.

**Fig. 6:**
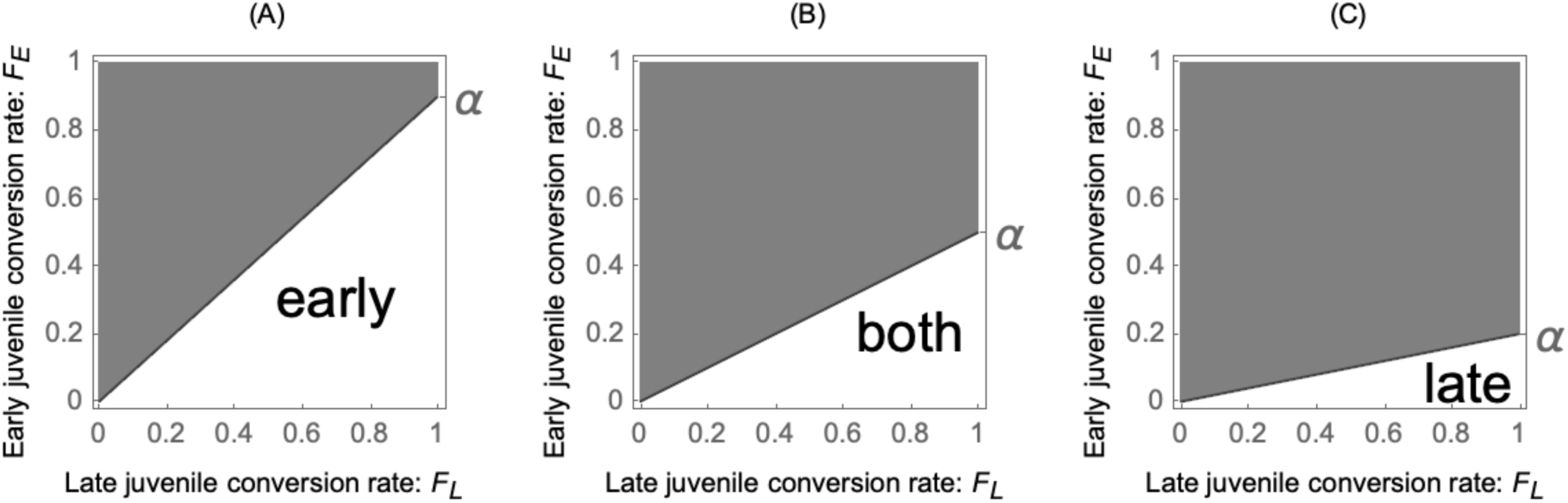
The evolutionary consequences of preference for host stages when delayed emergence occurs. Delayed emergence ( *y* = 1) is convergence stable in the white area, but not in the gray area. Parasitoids attack only early-stage hosts in the “early” area, attack both early and late-stage hosts in the “both” area, and attack late-stage hosts in the “late” area

#### B-6. Selection gradient of *y*

Selection gradient of *y* is given by:

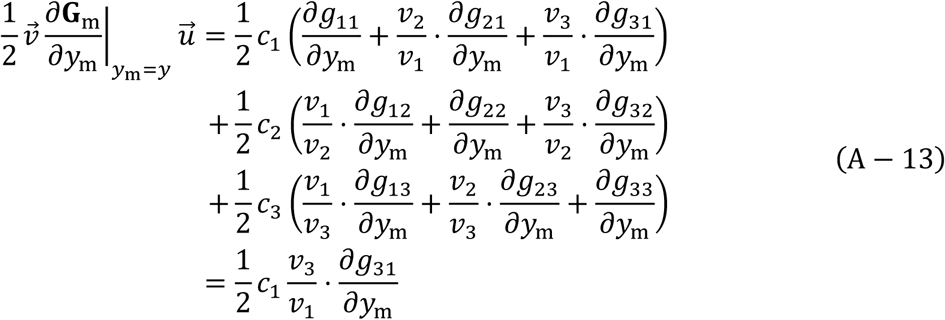

where

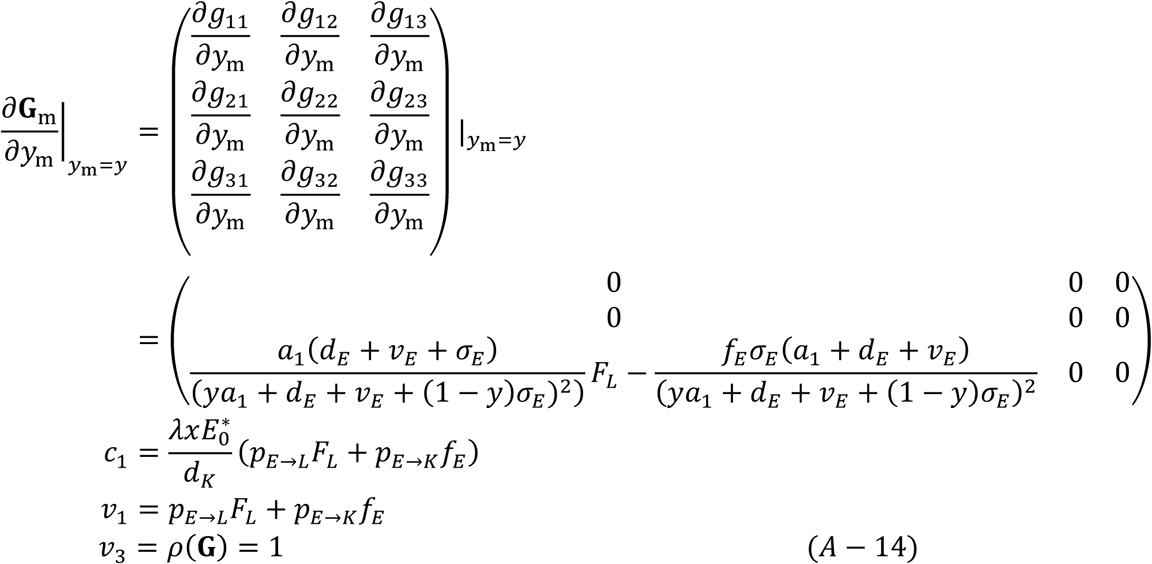

### C. Evolutionary stability

By examining the sign of the second derivative given that the candidate evolutionary consequence is in the interior of the phenotypic space (which is the square phenotypic space spanned by the two traits (*x*, *y*), we assess the evolutionary stability of *x* using the adaptive dynamics theory. We have now written as

The second derivative with respect to *x*_m_ is given by:

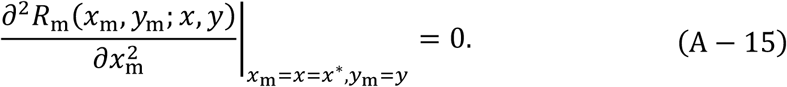

This follows from the equality 𝑤(*x*_𝑚_, *x*_𝑚_) = 𝑤(*x*^∗^, *x*_𝑚_). In this case, by definition, for the evolutionarily singular strategy (*x*^∗^) to be evolutionarily stable, the condition 𝑤(*x*^∗^, *x*) > 𝑤(*x*, *x*) must be satisfied.

For simplicity, let us express 𝑅_m_(*x*_m_, *y*_m_; *x*, *y*) as 𝑅_m_(*x*_m_, *x*) = *A*(*x*)*x*_m_ + 𝐵(*x*)(1 − *x*_m_). We guarantee the evolutionary stability by proving that 𝑅_m_(*x*_m_, *x*) > 𝑅_m_(x, *x*).

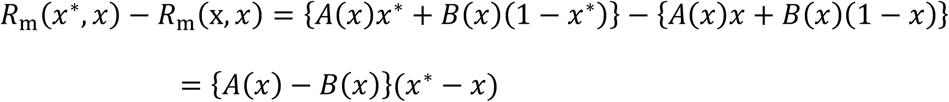

Here, equations (A-6) and (A-7) imply convergence stability. Since the selection gradient is written as *A*(*x*) − 𝐵(*x*),

If *x*^∗^ < *x*, *A*(*x*) − 𝐵(*x*) < 0

If *x*^∗^ > *x*, *A*(*x*) − 𝐵(*x*) > 0

Therefore, the inequality 𝑅_m_(*x*^∗^, *x*) − 𝑅_m_(x, *x*) > 0 always holds.

On the other hand, we can examine the evolutionary stability of the preference for delayed emergence *y* without taking the second derivative because the selection gradient of *y* is independent of frequency. The selection gradient of *y* is given by:

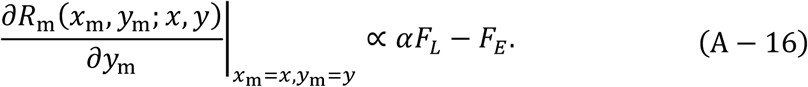

The selection pushes the trait to the boundary of the phenotypic space, causing evolution to stop there. Thus, when the sign is negative, *y*^∗^ = 0 is evolutionarily stable (i.e., the strategies that no other rare mutant with small phenotypic differences can increase in frequency) because it attains a local maximum. We can apply a similar argument to the case where *y*^∗^ = 1, and find that *y*^∗^ = 1 is also evolutionarily stable when the sign is positive, because it attains a local maximum.

### D. An unique equilibrium of the population dynamics

We can obtain the density ratio of *L*^∗^_0_ to *E*^∗^_0_ as follows:

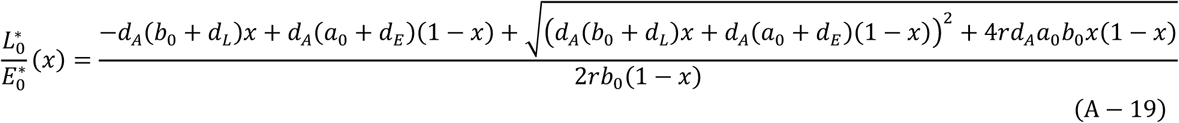

which is a monotone increasing function of *x*. Here, we will prove the uniqueness of the equilibrium of the population dynamics (Equation (1)). From the third line, we obtain

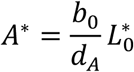

From the third line, we obtain *K*^∗^ analytically

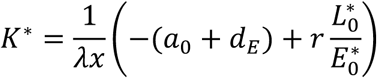

From the fourth line and the fifth line, we obtain

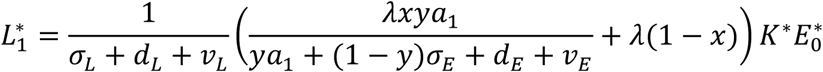

From the sixth line, we obtain

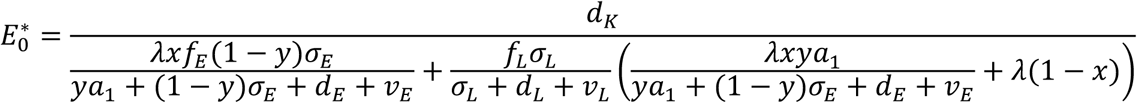

From this equation, we can obtain all the rest equilibrium *L*^∗^_0_, *A*^∗^, *E*_1_^∗^.

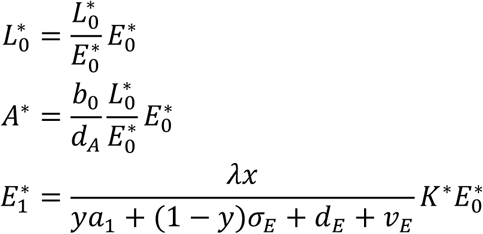

### E. Simulation

We conduct deterministic evolutionary simulations and then compare the results of exponential growth with those of logistic growth. First, we divide the parasitoids population into 25 strains based on the evolutionary traits and construct a mathematical model that represents the population dynamics of these strains. Second, we numerically solve the population dynamics to determine which strain survives over an extended period. Finally, we compare the outcomes of evolutionary simulations of the case where the host has logistic growth with that of the case where the host has exponential growth. We show that our analytical results are robust against this modified assumption.

**Fig. 7.**
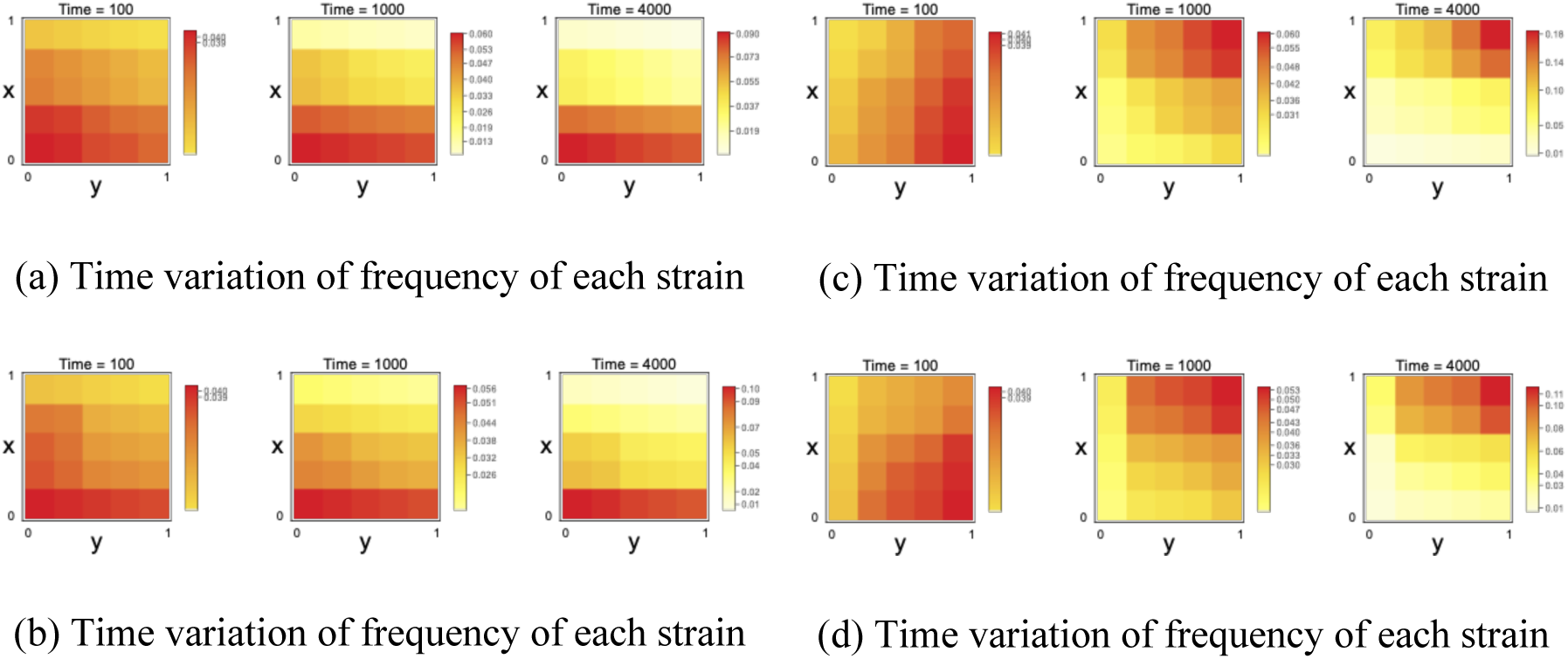
Evolutionary simulation: (a) The host density follows linear growth. (b) The host density follows logistic growth (carrying capacity 𝑐 = 100). Both represent the case where parasitoids evolve to idiobiont. Parasitoids attack late-stage hosts when *x* = 0, so the preference for delayed emergence *y* is neutral (*λ* = 1, *r* = 4, *a*_0_ = 1.5, *b*_0_ = 0.5, *d*_*E*_ = 1, *d*_*L*_ = 1, *d*_*A*_ = 0.5, *d*_*K*_ = 0.5, 𝑣_*E*_ = 0.5, 𝑣_*L*_ = 0.5, 𝜎_*E*_ = 0.5, 𝜎_*L*_ = 1, 𝑓_*E*_ = 0.5, 𝑓_*L*_ = 0.6). (c)The host density follows linear growth. (d)The host density follows logistic growth (The carrying capacity 𝑐=100). Both of Fig7 (c) and Fig7 (d) represent the case that parasitoids evolve to koinobiont (*λ* = 0.1, *r* = 0.5, *a*_0_ = 0.16, *b*_0_ = 0.2, *d*_*E*_ = 0.1, *d*_*L*_ = 0.1, *d*_*A*_ = 0.1, *d*_*K*_ = 0.1, 𝑣_*E*_ = 0.1, 𝑣_*L*_ = 0.1, 𝜎_*E*_ = 1, 𝜎_*L*_ = 1, 𝑓_*E*_ = 1, 𝑓_*L*_ = 2)

## Acknowledgement

We thank Troy Day, Koichi Ito, Bo-Moon Kim, Raiki Nakano, Stuart West, and Atsushi Yamauchi for useful comments.

## Author contributions

Ryuichiro Isshiki conceived the study, developed the theoretical framework, performed the mathematical analyses, and drafted the manuscript. Ryosuke Iritani provided supervision, critical feedback, and editorial revisions. All authors read and approved the final version of the manuscript.

